# Characterizing the genetic history of admixture across inner Eurasia

**DOI:** 10.1101/327122

**Authors:** Choongwon Jeong, Oleg Balanovsky, Elena Lukianova, Nurzhibek Kahbatkyzy, Pavel Flegontov, Valery Zaporozhchenko, Alexander Immel, Chuan-Chao Wang, Olzhas Ixan, Elmira Khussainova, Bakhytzhan Bekmanov, Victor Zaibert, Maria Lavryashina, Elvira Pocheshkhova, Yuldash Yusupov, Anastasiya Agdzhoyan, Koshel Sergey, Andrei Bukin, Pagbajabyn Nymadawa, Michail Churnosov, Roza Skhalyakho, Denis Daragan, Yuri Bogunov, Anna Bogunova, Alexandr Shtrunov, Nadezda Dubova, Maxat Zhabagin, Levon Yepiskoposyan, Vladimir Churakov, Nikolay Pislegin, Larissa Damba, Ludmila Saroyants, Khadizhat Dibirova, Lubov Artamentova, Olga Utevska, Eldar Idrisov, Evgeniya Kamenshchikova, Irina Evseeva, Mait Metspalu, Martine Robbeets, Leyla Djansugurova, Elena Balanovska, Stephan Schiffels, Wolfgang Haak, David Reich, Johannes Krause

## Abstract

The indigenous populations of inner Eurasia, a huge geographic region covering the central Eurasian steppe and the northern Eurasian taiga and tundra, harbor tremendous diversity in their genes, cultures and languages. In this study, we report novel genome-wide data for 763 individuals from Armenia, Georgia, Kazakhstan, Moldova, Mongolia, Russia, Tajikistan, Ukraine, and Uzbekistan. We furthermore report genome-wide data of two Eneolithic individuals (∽5,400 years before present) associated with the Botai culture in northern Kazakhstan. We find that inner Eurasian populations are structured into three distinct admixture clines stretching between various western and eastern Eurasian ancestries. This genetic separation is well mirrored by geography. The ancient Botai genomes suggest yet another layer of admixture in inner Eurasia that involves Mesolithic hunter-gatherers in Europe, the Upper Paleolithic southern Siberians and East Asians. Admixture modeling of ancient and modern populations suggests an overwriting of this ancient structure in the Altai-Sayan region by migrations of western steppe herders, but partial retaining of this ancient North Eurasian-related cline further to the North. Finally, the genetic structure of Caucasus populations highlights a role of the Caucasus Mountains as a barrier to gene flow and suggests a post-Neolithic gene flow into North Caucasus populations from the steppe.

## Introduction

Present-day human population structure is often marked by a correlation between geographic and genetic distances,^1; 2^ reflecting continuous gene flow among neighboring groups, a process known as “isolation by distance”. However, there are also striking failures of this model, whereby geographically proximate populations can be quite distantly related. Such barriers to gene flow often correspond to major geographic features, such as the Himalayas^3^ or the Caucasus Mountains.^4^ Many cases also suggest the presence of social barriers to gene flow. For example, early Neolithic farming populations in Europe show a remarkable genetic homogeneity suggesting minimal genetic exchange with local hunter-gatherer populations through the initial expansion; genetic mixing of these two gene pools became evident only after thousands of years in the middle Neolithic.^5^ Modern Lebanese populations provide another example by showing a population stratification reflecting their religious community.^6^ There are also examples of geographically very distant populations that are closely related: for example, people buried in association with artifacts of the Yamnaya horizon in the Pontic-Caspian steppe and the contemporaneous Afanasievo culture 3,000 km east in the Altai-Sayan Mountains.^7; 8^

The vast region of the Eurasian inland (“inner Eurasia” herein) is split into distinct ecoregions, such as the Eurasian steppe in central Eurasia, boreal forests (taiga) in northern Eurasia, and the Arctic tundra at the periphery of the Arctic Ocean. These ecoregions stretch in an east-west direction within relatively narrow north-south bands. Various cultural features show a distribution that broadly mirrors the eco-geographic distinction in inner Eurasia. For example, indigenous peoples of the Eurasian steppe traditionally practice nomadic pastoralism,^9; 10^ while northern Eurasian peoples in the taiga mainly rely on reindeer herding and hunting^11^. The subsistence strategies in each of these ecoregions are often considered to be adaptations to the local environments.^12^

At present there is limited information about how environmental and cultural influences are mirrored in the genetic structure of inner Eurasians. Recent genome-wide studies of inner Eurasians mostly focused on detecting and dating genetic admixture in individual populations.^13-16^ So far only two studies have reported recent genetic sharing between geographically distant populations based on the analysis of “identity-by-descent” segments.^13; 17^ One study reports a long-distance extra genetic sharing between Turkic populations based on a detailed comparison between Turkic-speaking groups and their non-Turkic neighbors.^13^ Another study extends this approach to some Uralic and Yeniseian-speaking populations.^17^ However, a comprehensive spatial genetic analysis of inner Eurasian populations is still lacking.

Ancient DNA studies have already shown that human populations of this region have dramatically transformed over time. For example, the Upper Paleolithic genomes from the Mal’ta and Afontova Gora archaeological sites in southern Siberia revealed a genetic profile, often referred to as “Ancient North Eurasians (ANE)”, which is deeply related to Paleolithic/Mesolithic hunter-gatherers in Europe and also substantially contributed to the gene pools of modern-day Native Americans, Siberians, Europeans and South Asians.^18; 19^ Studies of Bronze Age steppe populations found the appearance of additional Western Eurasian-related ancestries across the steppe from the Pontic-Caspian region in the West to the Altai-Sayan region in the East, here we collectively refer to as “Western Steppe Herders (WSH)”: the earlier populations associated with the Yamnaya and Afanasievo cultures (often referred to “steppe Early and Middle Bronze Age”; “steppe_EMBA”) and the later ones associated with many cultures such as Potapovka, Sintashta, Srubnaya and Andronovo to name a few (often referred to “steppe Middle and Late Bronze Age”; “steppe_MLBA”).^8^ Still, important questions remain unanswered due to limited availability of ancient genomes, including the identity of the eastern Eurasian gene pools that interacted with Pleistocene ANE or Bronze Age WSH populations and the genetic profile of pre-Bronze Age inner Eurasians. An example of the latter is the Eneolithic Botai culture in northern Kazakhstan in the 4^th^ millennium BCE.^20^ In addition to their role in the earliest horse domestication so far known,^21^ Botai is at the crossroads, both in time and in space, connecting various earlier hunter-gatherer and later WSH populations in inner Eurasia.

In this study, we analyzed newly produced genome-wide genetic variation data for 763 individuals belonging to 60 self-reported ethnic groups to provide a dense portrait of the genetic structure of indigenous populations in inner Eurasia. We also produced genome-wide data of two individuals associated with the Eneolithic Botai culture in Kazakhstan to explore the genetic structure of pre-Bronze Age populations in inner Eurasia. We aimed at characterizing the genetic composition of inner Eurasians in fine resolution by applying both allele frequency- and haplotype-based methods. Based on the fine-scale genetic profile, we further explored if and where the barriers and conduits of gene flow exist in inner Eurasia.

## Materials and Methods

### Study participants and genotyping

We collected samples from 763 participants from nine countries (Armenia, Georgia, Kazakhstan, Moldova, Mongolia, Russia, Tajikistan, Ukraine, and Uzbekistan). The sampling strategy included sampling a majority of large ethnic groups in the studied countries. Within groups, we sampled subgroups if they were known to speak different dialects; for ethnic groups with large area, we sampled within several districts across the area. We sampled individuals whose grandparents were all self-identified members of the given ethnic groups and were born within the studied district(s). All individuals provided a written informed consent approved by the Ethic Committee of the Research Centre for Medical Genetics, Moscow, Russia. Most of the ethnic Russian samples were collected from indigenous Russian areas (present-day Central Russia) and had been stored for years in the Estonian Biocenter; samples from Mongolia, Tajikistan, Uzbekistan, and Ukraine were collected partially in the framework of the Genographic project. Most DNA samples were extracted from venous blood via the phenol-chloroform method. For this study we identified 112 subgroups (belonging to 60 ethnic group labels) which were not previously genotyped on the Affymetrix Axiom^®^ Genome-wide Human Origins 1 (“HumanOrigins”) array platform^22^ and selected on average 7 individuals per subgroup (Figure 1 and Table S1). Genome-wide genotyping experiments were performed on the HumanOrigins array platform. We removed 18 individuals from further analysis either due to high genotype missing rate (> 0.05; n=2) or due to being outliers in principal component analysis (PCA) relative to other individuals from the same group (n = 16). The remaining 745 individuals assigned to 60 group labels were merged to published HumanOrigins data sets of world-wide contemporary populations^19^ and of four Siberian ethnic groups (Enets, Kets, Nganasans and Selkups).^23^ Diploid genotype data of six contemporary individuals (two Saami, two Sherpa and two Tibetans) were obtained from the Simons Genome Diversity Panel data set.^24^ We also added ancient individuals from published studies,^3; 8; 18; 19; 25-40^ by randomly sampling a single allele for 581,230 autosomal single nucleotide polymorphisms (SNPs) in the HumanOrigins array (Table S2).

**Figure 1.**
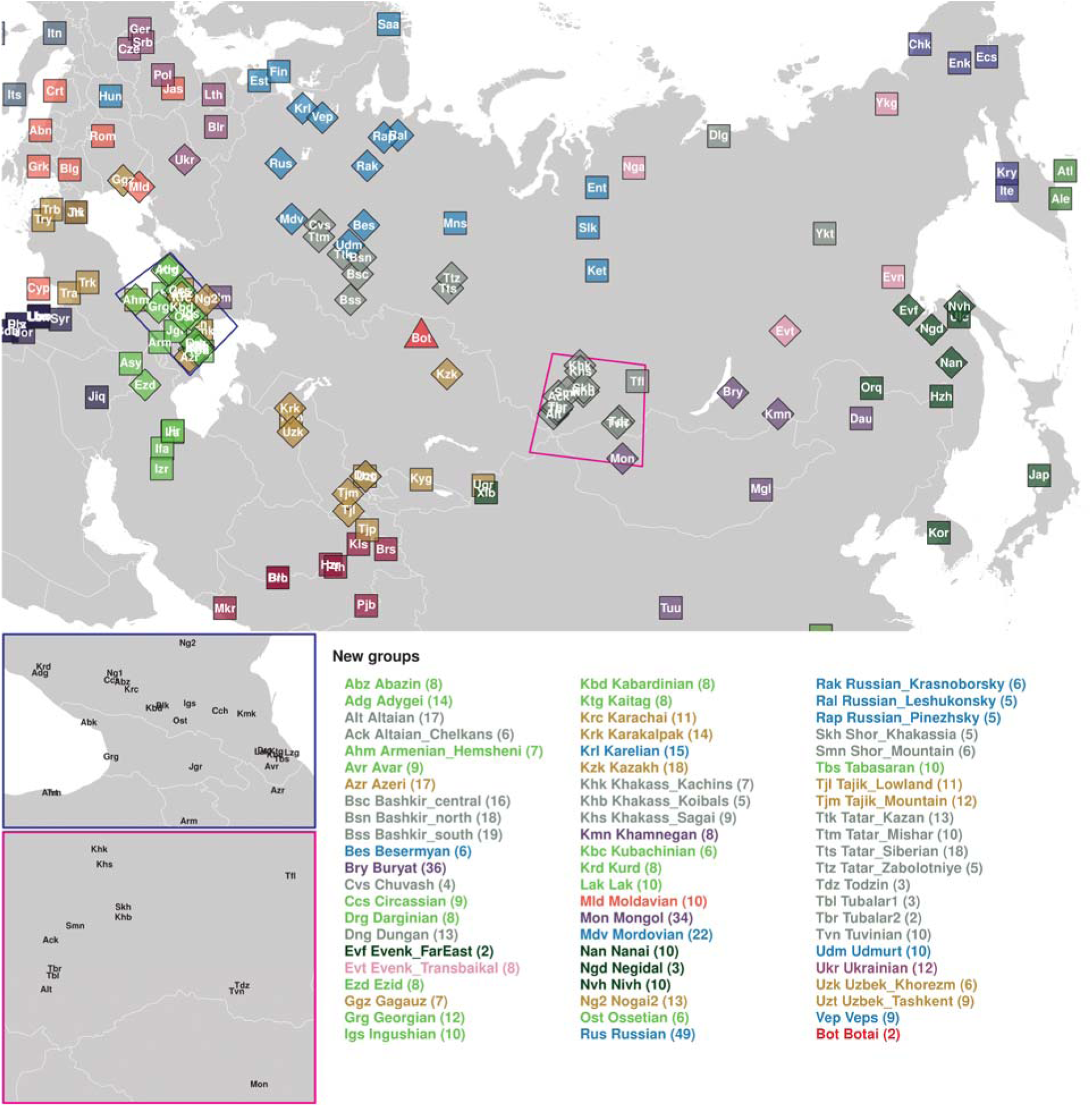
Geographic locations of the Eneolithic Botai site (red triangle), 65 groups including newly sampled individuals (filled diamonds) and nearby groups with published data (filled squares). Mean latitude and longitude values across all individuals under each group label were used. Two zoom-in plots for the Caucasus (blue) and the Altai-Sayan (magenta) regions are presented in the lower left corner. A list of new groups, their three-letter codes, and the number of new individuals (in parenthesis) are provided at the bottom. Corresponding information for the previously published groups is provided in Table S2. The main inner Eurasian map is on the Albers equal area projection and was produced using the spTransform function in the R package rgdal v1.2-5.

### Sequencing of the ancient Botai genomes

We extracted genomic DNA from four skeletal remains belonging to two individuals and built sequencing libraries either with no uracil-DNA glycosylase (UDG) treatment or with partial treatment following published protocols (Table 1).^41; 42^ Radiocarbon dating of BKZ001 was conducted by the CEZ Archaeometry gGmbH (Mannheim, Germany) for one of two bone samples used for DNA extraction. All libraries were barcoded with two library-specific 8-mer indices.^43^ The samples were manipulated in dedicated clean room facilities at the University of Tübingen or at the Max Planck Institute for the Science of Human History (MPI-SHH). Indexed libraries were enriched for about 1.24 million informative nuclear SNPs using the in-solution capture method (“1240K capture”).^5; 31^

**Table 1.** Sequencing statistics and radiocarbon dates of two Eneolithic Botai individuals analyzed in this study.

| ID | Genetic Sex | Uncal.<br><sup>14</sup>C Date | Cal.<br><sup>14</sup>C Date<br>(2-sigma)<sup>b</sup> | # of reads<br>sequenced | # of SNPs<br>covered<sup>c</sup> | MT / Y<br>haplogroup | MT.cont<sup>d</sup> | X.cont<sup>e</sup> |
| --- | --- | --- | --- | --- | --- | --- | --- | --- |
| TU45 | M | 4620<br>± 80<sup>a</sup> | 3632-3100<br>cal. BCE | 84,170,835 | 77,363 | K1b2 /<br>R1b1a1 | 0.02<br>(0.01-0.03) | 0.0122<br>(0.0050) |
| BKZ001 | F | 4660<br>± 25 | 3517-3367<br>cal. BCE | 69,678,735 | 432,078 | Z1 / NA | 0.01<br>(0.00-0.02) | NA |

<sup>a</sup> The uncalibrated date of TU45 was published in Levine (1999) under the ID OxA-4316.<sup>70</sup>
<sup>b</sup> The calibrated <sup>14</sup>C dates are calculated based on uncalibrated dates, by the OxCal v4.3.2 program<sup>71</sup> using the INTCAL13 atmospheric curve.<sup>72</sup>
<sup>c</sup> The number of autosomal SNPs in the HumanOrigins array (out of 581,230) covered at least by one read. Only transversion SNPs are considered for the non-UDG libraries (both of the TU45 libraries, one of two BKZ001 libraries).
<sup>d</sup> The contamination rate of mitochondrial reads estimated by the Schmutzi program (95% confidence interval in parentheses)
<sup>e</sup> The nuclear contamination rate for the male (TU45) estimated based on X chromosome data by ANGSD software (standard error in parentheses)

Libraries were sequenced on the Illumina HiSeq 4000 platform with either single-end 75 bp (SE75) or paired-end 50 bp (PE50) cycles following manufacturer’s protocols. Output reads were demultiplexed by allowing up to 1 mismatch in each of two 8-mer indices. FASTQ files were processed using EAGER v1.92.^44^ Specifically, Illumina adapter sequences were trimmed using AdapterRemoval v2.2.0,^45^ aligned reads (30 base pairs or longer) onto the human reference genome (hg19) using BWA aln/samse v0.7.12^46^ with relaxed edit distance parameter (“-n 0.01”). Seeding was disabled for reads from non-UDG libraries by adding an additional parameter (“-l 9999”). PCR duplicates were then removed using DeDup v0.12.2^44^ and reads with Phred-scaled mapping quality score < 30 were filtered out using Samtools v1.3.^47^ We did several measurements to check data authenticity. First, patterns of chemical damages typical to ancient DNA were tabulated using mapDamage v2.0.6.^48^ Second, mitochondrial contamination for all libraries was estimated by Schmutzi.^49^ Third, nuclear contamination for libraries derived from males was estimated by the contamination module in ANGSD v0.910.^50^ Prior to genotyping, the first and last 3 bases of each read were masked for libraries with partial UDG treatment using the trimBam module in bamUtil v1.0.13.^51^ To obtain haploid genotypes, we randomly chose one high-quality base (Phred-scaled base quality score ≥ 30) for each of the 1.24 million target sites using pileupCaller (https://github.com/stschiff/sequenceTools). We used masked reads from libraries with partial UDG treatment for transition (Ts) SNPs and used unmasked reads from all libraries for transversions (Tv). Mitochondrial consensus sequences were obtained by the log2fasta program in Schmutzi with the quality cutoff 10 and subsequently assigned to haplogroups using HaploGrep2.^52^ Y haplogroup R1b was assigned using the yHaplo program.^53^ To estimate the phylogenetic position of the Botai Y haplogroup more precisely, Y chromosomal SNPs were called with Samtools mpileup using bases with quality score ≥ 30: a total of 2,481 SNPs out of ∽30,000 markers included in the 1240K capture panel were called with mean read depth of 1.2. Twenty-two SNP positions relevant to the up-to-date haplogroup R1b tree (www.isogg.org; www.yfull.com) confirmed that the sample was positive for the markers of R1b-P297 branch but negative for its R1b-M269 sub-branch.

The frequency distribution map of this Y chromosomal clade was created by the GeneGeo software^54; 55^ using the average weighed interpolation procedure with the weight function of degree 3 and radius 1, 200 km. The initial frequencies were calculated as proportion of samples positive for “root” R1b marker M343 but negative for M269; these proportions were calculated for the 577 populations from the in-home *Y-base* database, which was compiled mainly from the published datasets.

### Analysis of population structure

We performed principal component analysis (PCA) of various groups using smartpca v13050 in the EIGENSOFT v6.0.1 package.^56^ We used the “*lsqproject: YES*” option to project individuals not used for calculating PCs (this procedure avoids bias due to missing genotypes). We performed unsupervised model-based genetic clustering as implemented in ADMIXTURE v1.3.0.^57^ For that purpose, we used 116,468 SNPs with minor allele frequency (maf) 1% or higher in 3,332 individuals after pruning out linked SNPs (r^2^ > 0.2) using the “--indep-pairwise 200 25 0.2” command in PLINK v1.90.^58^ We then converted diploid genotypes to haploid data by randomly choosing one of the two alleles to minimize a bias due to artificial genetic drift in haploid genotype calls of most low coverage ancient individuals. For each value of K ranging from 2 to 20, we ran 5 replicates with different random seeds and took one with the highest log likelihood value.

### F-statistics analysis

We computed various *f*_*3*_ and *f*_*4*_ statistics using the qp3Pop (v400) and qpDstat (v711) programs in the ADMIXTOOLS package.^22^ We computed *f*_*4*_-statistics with the “*f4mode: YES*” option. For these analyses, we studied a total of 301 groups, including 73 inner Eurasian target groups and 167 contemporary and 61 ancient reference groups (Table S2). We included two groups from the Aleutian Islands (“Aleut” and “Aleut_Tlingit”; Table S2) as positive control targets with known recent admixture. Aleut_Tlingits are Aleut individuals whose mitochondrial haplogroup lineages are related to Tlingits.^29^ For each target, we calculated outgroup *f*_*3*_ statistic of the form *f*_*3*_(Target, X; Mbuti) against all targets and references to quantify overall allele sharing and performed admixture *f*_*3*_ test of the form *f*_*3*_(Ref_1_, Ref_2_; Target) for all pairs of references to explore the admixture signal in targets. We estimated standard error (SE) using a block jackknife with 5 centiMorgan (cM) block.^56^

We performed *f*_*4*_ statistic-based admixture modeling using the qpAdm (v632) program^19^ in the ADMIXTOOLS package. We used a basic set of 7 outgroups, unless specified otherwise, to provide high enough resolution to distinguish various western and eastern Eurasian ancestries: Mbuti (n=10; central African), Natufian (n=6; early Holocene Levantine),^19^ Onge (n=11; from the Andaman Islands), Iran_N (n=5; Neolithic Iranian),^19^ Villabruna (n=1; Paleolithic European),^26^ Ami (n=10; Taiwanese aborigine) and Mixe (n=10; Central American). Prior to qpAdm modeling, we checked if the reference groups are well distinguished by their relationship with the outgroups using the qpWave (v400) program.^59^

We used the qpGraph (v6065) program in the ADMIXTOOLS package for graph-based admixture modeling. Starting with a graph of (Mbuti, Ami, WHG), we iteratively added AG3 (n=1; Paleolithic Siberian),^26^ EHG (n=3; Mesolithic hunter-gatherers from Karelia or Samara),^5; 26^ and Botai onto the graph by testing all possible topologies allowing up to one additional gene flow. After obtaining the best two-way admixture model for Botai, we tested additional three-way admixture models.

### GLOBETROTTER analysis

We performed a GLOBETROTTER analysis of admixture for 73 inner Eurasian target populations to obtain haplotype sharing based evidence of admixture, independent of the allele frequency based *f*-statistics, as well as estimates of admixture dates and a fine-scale profile of their admixture sources.^14^ We followed the “regional” approach described in Hellenthal et al.,^14^ in which target haplotypes can only be copied from the haplotypes of 167 contemporary reference groups, but not from those of the other target groups. This approach is recommended when multiple target groups share a similar admixture history,^14^ which is likely to be the case for our inner Eurasian populations.

We jointly phased the contemporary genome data without a pre-phased set of reference haplotypes, using SHAPEIT2 v2.837 in its default setting.^60^ We used a genetic map for the 1000 Genomes Project phase 3 data, downloaded from: https://mathgen.stats.ox.ac.uk/impute/1000GP_Phase3.html. We used haplotypes from a total of 2,615 individuals belonging to 240 groups (73 recipients and 167 donors; Table S2) for the GLOBETROTTER analysis. To reduce computational burden and to provide more balanced set of donor populations, we randomly sampled 20 individuals if a group contained more than 20 individuals. Using these haplotypes, we performed GLOBETROTTER analysis following the recommended workflow.^14^ We first ran 10 rounds of the expectation-maximization (EM) algorithm for chromosomes 4, 10, 15 and 22 in ChromoPainter v2 with “-in” and “-iM” switches to estimate chunk size and switch error rate parameters.^61^ Both recipient and donor haplotypes were modeled as a patchwork of donor haplotypes. The “chunk length” output was obtained by running ChromoPainter v2 across all chromosomes with the estimated parameters averaged over both recipient and donor individuals (“-n 238.05 -M 0.000617341”). We also generated 10 painting samples for each recipient group by running ChromoPainter with the parameters averaged over all recipient individuals (“-n 248.455 -M 0.000535236”). Using the chunklength output and painting samples, we ran GLOBETROTTER with the “prop.ind: 1” and “null.ind: 1” options. We estimated significance of estimated admixture date by running 100 bootstrap replicates using the “prop.ind: 0” and “bootstrap.date.ind: 1” options; we considered date estimates between 1 and 400 generations as evidence of admixture.^14^ For populations that gave evidence of admixture by this procedure, we repeated GLOBETROTTER analysis with the “null:ind: 0” option.^14^ We also compared admixture dates from GLOBETROTTER analysis with those based on weighted admixture linkage disequilibrium (LD) decay, as implemented in ALDER v1.3.^62^ As the reference pair, we used (French, Eskimo_Naukan), (French, Nganasan), (Georgian, Ulchi), (French, Ulchi) and (Georgian, Ulchi) for the target group categories 1 to 5, respectively, based on their genetic profile (Table S2). We used a minimum inter-marker distance of 1.0 cM to account for LD in the references.

### EEMS analysis

To visualize the heterogeneity in the rate of gene flow across inner Eurasia, we performed the EEMS (“estimated effective migration surface”) analysis.^63^ We included a total of 1,180 individuals from 94 groups in the analysis (Table S2). In this dataset, we kept 101,320 SNPs with maf ≥ 0.01 after LD pruning (r^2^ ≤ 0.2). We computed the mean squared genetic difference matrix between all pairs of individuals using the “bed2diffs_v1” program in the EEMS package. To reduce distortion in northern latitudes due to map projection, we used geographic coordinates in the Albers equal area conic projection (“+ proj=aea + lat_1=50 + lat_2=70 + lat_0=56 + lon_0=100 + x_0=0 + y_0=0 + ellps=WGS84 + datum=WGS84 + units=m + no_defs”). We converted geographic coordinates of each sample and the boundary using the “spTransform” function in the R package rgdal v1.2-5. We ran five initial MCMC runs of 2 million burn-ins and 4 million iterations with different random seeds and took a run with the highest likelihood. Starting from the best initial run, we set up another five MCMC runs of 2 million burn-ins and 4 million iterations as our final analysis. We used the following proposal variance parameters to keep the acceptance rate around 30-40%, as recommended by the developers^63^: qSeedsProposalS2 = 5000, mSeedsProposalS2 = 1000, qEffctProposalS2 = 0.0001, mrateMuProposalS2 = 0.00005. We set up a total of 532 demes automatically with the “nDemes = 600” parameter. We visualized the merged output from all five runs using the “eems.plots” function in the R package rEEMSplots.^63^

We performed the EEMS analysis for Caucasus populations in a similar manner, including a total of 237 individuals from 21 groups (Table S2). In this dataset, we kept 95,442 SNPs with maf ≥ 0.01 after LD pruning (r^2^ ≤ 0.2). We applied the Mercator projection of geographic coordinates to the map of Eurasia (“+ proj=merc + datum=WGS84”). We ran five initial MCMC runs of 2 million burn-ins and 4 million iterations with different random seeds and took a run with the highest likelihood. Starting from the best initial run, we set up another five MCMC runs of 1 million burn-in and 4 million iterations as our final analysis. We used the default following proposal variance parameters: qSeedsProposalS2 = 0.1, mSeedsProposalS2 = 0.01, qEffctProposalS2 = 0.001, mrateMuProposalS2 = 0.01. A total of 171 demes were automatically set up with the “nDemes = 200” parameter.

## Results

### Inner Eurasians form distinct east-west genetic clines mirroring geography

In a PCA of Eurasian individuals, we find that PC1 separates eastern and western Eurasian populations, PC2 splits eastern Eurasians along a north-south cline, and PC3 captures variation in western Eurasians with Caucasus and northeastern European populations at opposite ends (Figure 2A and Figures S1-S2). Inner Eurasians are scattered across PC1 in between, largely reflecting their geographic locations. Strikingly, inner Eurasian populations seem to be structured into three distinct west-east genetic clines running between different western and eastern Eurasian groups, instead of being evenly spaced in PC space. Individuals from northern Eurasia, speaking Uralic or Yeniseian languages, form a cline connecting northeast Europeans and the Uralic (Samoyedic) speaking Nganasans from northern Siberia (“forest-tundra” cline). Individuals from the Eurasian steppe, mostly speaking Turkic and Mongolic languages, are scattered along two clines below the forest-tundra cline. Both clines run into Turkic- and Mongolic-speaking populations in southern Siberia and Mongolia, and further into Tungusic-speaking populations in Manchuria and the Russian Far East in the East; however, they diverge in the west, one heading to the Caucasus and the other heading to populations of the Volga-Ural area (the “southern steppe” and “steppe-forest” clines, respectively; Figure 2 and Figure S2).

**Figure 2.**
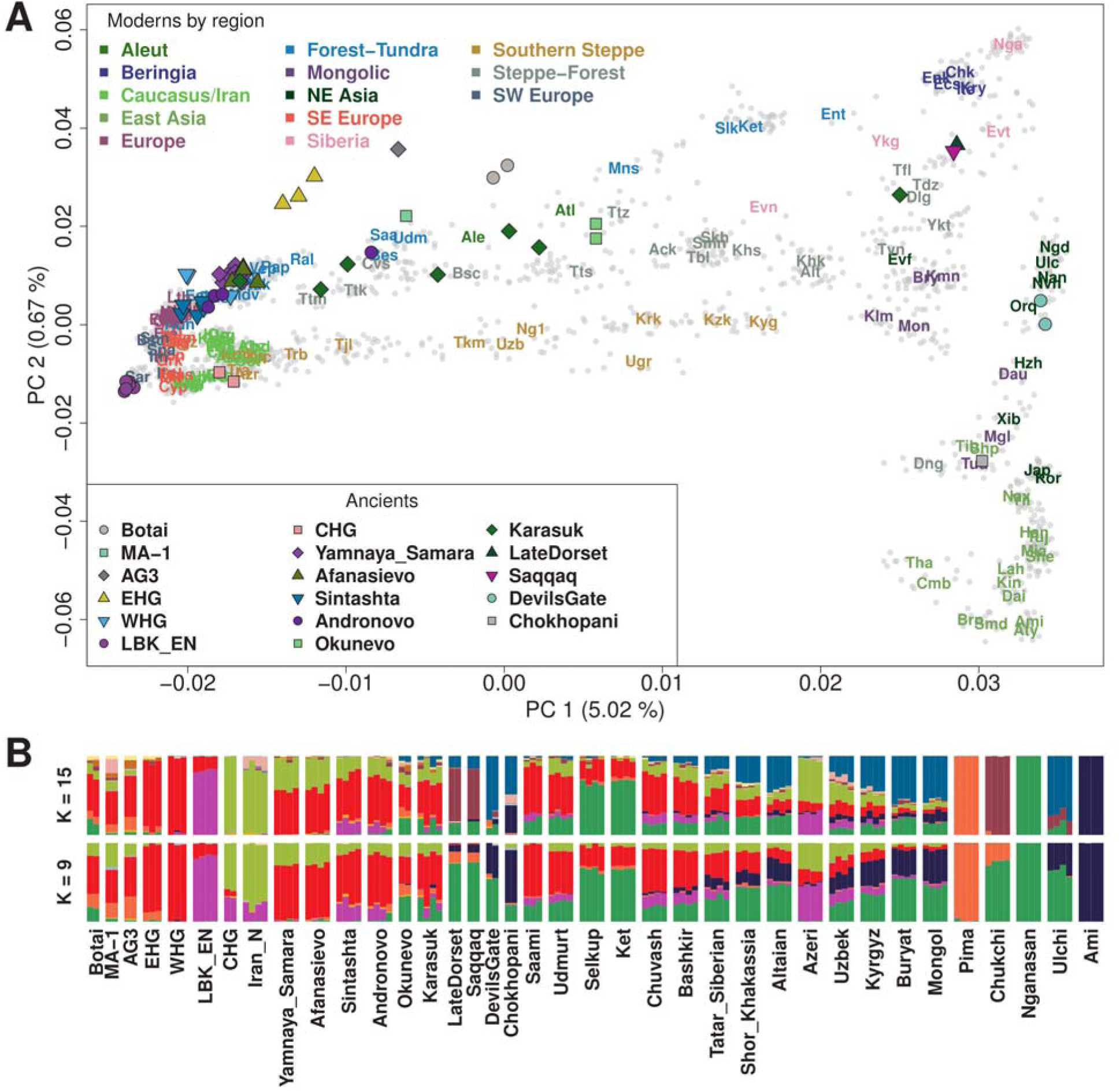
The genetic structure of inner Eurasian populations. (A) The first two PCs of 2,077 Eurasian individuals separate western and eastern Eurasians (PC1) and Northeast and Southeast Asians (PC2). Most inner Eurasians are located between western and eastern Eurasians on PC1. Ancient individuals (color-filled shapes) are projected onto PCs calculated based on contemporary individuals. Modern individuals are marked by grey dots, with their per-group mean coordinates marked by three-letter codes listed in Table S2. (B) ADMIXTURE results for a chosen set of ancient and modern groups (K = 9 and 15). Most inner Eurasians are modeled as a mixture of components primarily found in eastern or western Eurasians. Results for the full set of individuals are provided in Figure S3.

A model-based clustering analysis using ADMIXTURE shows a similar pattern (Figure 2B and Figure S3). Overall, the proportions of ancestry components associated with eastern or western Eurasians are well correlated with longitude in inner Eurasians (Figure 3A). Notable outliers from this trend include known historical migrants such as Kalmyks, Nogais and Dungans. The forest-tundra cline populations derive most of their eastern Eurasian ancestry from a component most enriched in Nganasans, while those on the steppe-forest and southern steppe clines have this component together with another component most enriched in populations from the Russian Far East, such as Ulchi and Nivkh. The southern steppe cline groups are distinct from the others in their western Eurasian ancestry profile, in the sense that they have a high proportion of a component most enriched in Mesolithic Caucasus hunter-gatherers (“CHG”)^28^ and Neolithic Iranians (“Iran_N”)^19^ and frequently harbor another component enriched in South Asians (Figure S4).

**Figure 3.**
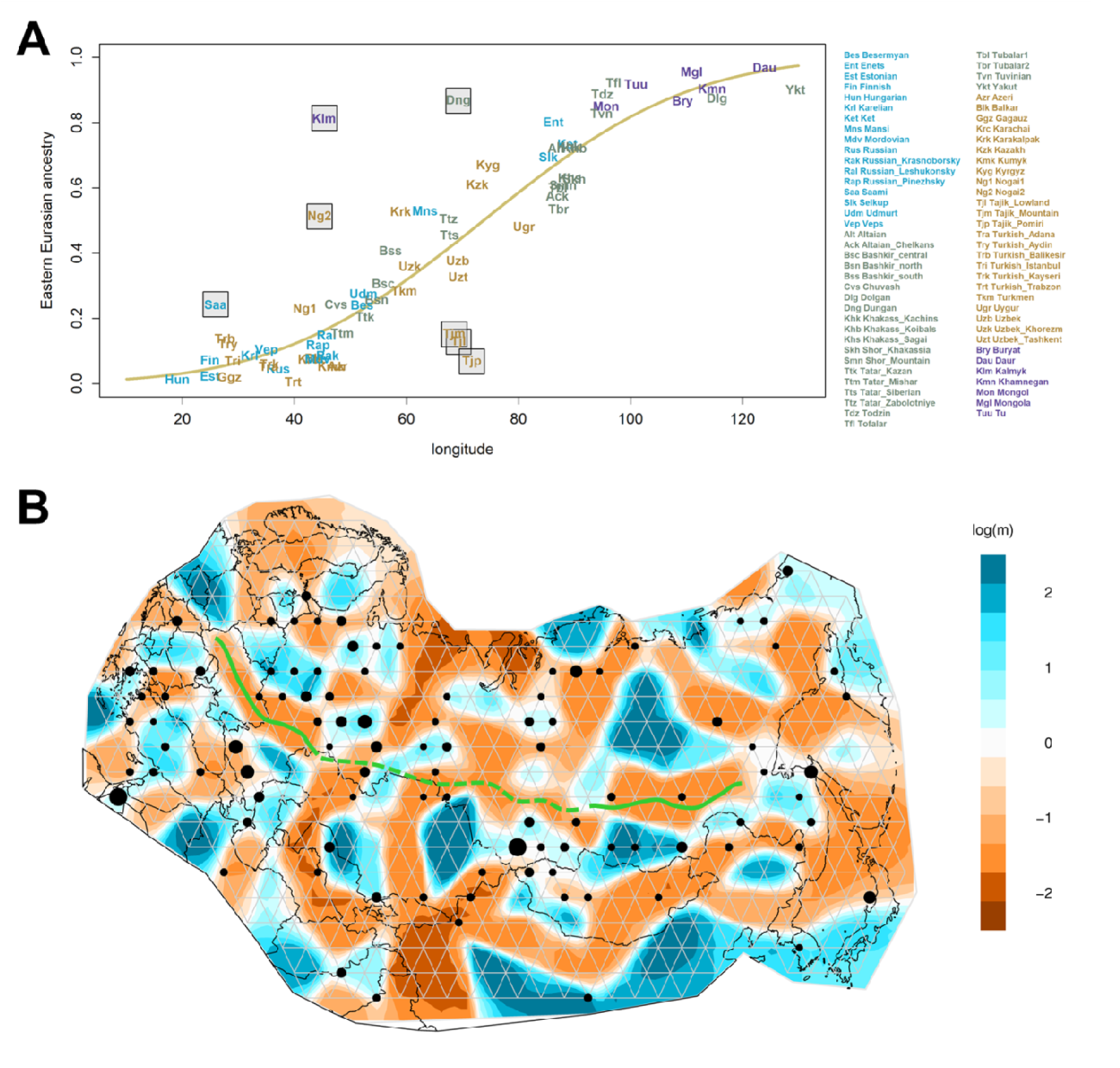
Inner Eurasian admixture in geographical context. (A) A comparison of mean longitudinal coordinates (x-axis) and mean eastern Eurasian ancestry proportions (y-axis) of inner Eurasians. Eastern Eurasian ancestry proportions are estimated from ADMIXTURE results with K = 15 by summing up six components maximized in Karitiana, Pima, Chukchi, Nganasan, Ulchi and Ami, respectively (Figure S3). The yellow curve shows a probit regression fit following the model in Sedghifar et al.^69^ Seven groups substantially deviating from the curve, including known historical migrants, are marked with grey background. (B) Barriers (brown) and conduits (blue) of gene flow across inner Eurasia estimated by the EEMS program. Black dots show the location of vertices to which individuals are assigned, with sizes correlated with the number of individuals. Solid green curves highlight strong barriers to gene flow separating the steppe-forest cline and the southern steppe cline populations (the western curve) or the steppe-forest cline and the forest-tundra cline populations (the eastern curve). The dotted green curve marks a region between the two curves where this barrier seems to be weaker than in the flanking regions.

The genetic barriers splitting the inner Eurasian clines are also evidenced in the EEMS (“estimated effective migration surface”) analysis (Figure 3B). A strong genetic barrier is detected between the Caucasus and the Pontic-Caspian steppe regions, separating the southern steppe and steppe-forest clines. On the eastern side, another barrier north of Lake Baikal separates southern Siberians from the forest-tundra cline groups in the North. These two barriers are partially connected by a weaker barrier north of the Altai-Sayan region, likely reflecting both the east-west connection within the steppe-forest cline and the north-south connection along the Yenisei River.

### High-resolution tests of admixture distinguish the genetic profile of source populations in the inner Eurasian clines

We performed both allele frequency-based three-population (*f*_*3*_) tests and a haplotype-sharing-based GLOBETROTTER analysis to characterize the admixed gene pools of inner Eurasian groups. For these group-based analyses, we manually removed 87 outliers from our contemporary individuals based on PCA results (Table S1). We also split a few inner Eurasian groups showing genetic heterogeneity into subgroups based on PCA results and their sampling locations (Table S1). This was done to minimize false positive admixture signals. We chose 73 groups as the targets of admixture tests and another 228 groups (167 contemporary and 61 ancient groups) as the “sources” to represent world-wide genetic diversity (Table S2).

Testing all possible pairs of 167 contemporary “source” groups as references, we detect highly significant *f*_*3*_ statistics for 66 of 73 targets (< −3 SE; standard error; Table S3). Negative *f*_*3*_ values mean that allele frequencies of the target group are on average intermediate between the allele frequencies of the reference populations, providing unambiguous evidence that the target population is a mixture of groups related, perhaps deeply, to the source populations.^22^ Extending the references to include 61 ancient groups, we find that the seven non-significant groups also have small *f*_*3*_ statistics around zero (−5.1 SE to + 2.7 SE). Reference pairs with the most negative *f*_*3*_ statistics for the most part involve one eastern Eurasian and one western Eurasian group supporting the qualitative impression of east-west admixture from PCA and ADMIXTURE analysis. To highlight the difference between the distinct inner Eurasian clines, we looked into *f*_*3*_ results with representative reference pairs comprising two western Eurasian (French to represent Europeans and Georgian to represent Caucasus populations) and three eastern Eurasian groups (Nganasan, Ulchi and Korean). In the populations of the southern steppe cline, reference pairs with Georgians tend to produce more negative *f*_*3*_ statistics than those with French while the opposite pattern is observed for the steppe-forest and forest-tundra populations (Figure 4A). Reference pairs with Nganasans mostly result in more negative *f*_*3*_ statistic than those with Ulchi in the forest-tundra populations, but the opposite pattern is dominant in the southern steppe populations. Populations of the steppe-forest cline show an intermediate pattern: the northern ones tend to have more negative *f*_*3*_ statistics with Nganasans while the southern ones tend to have more negative *f*_*3*_ statistics with Ulchi.

**Figure 4.**
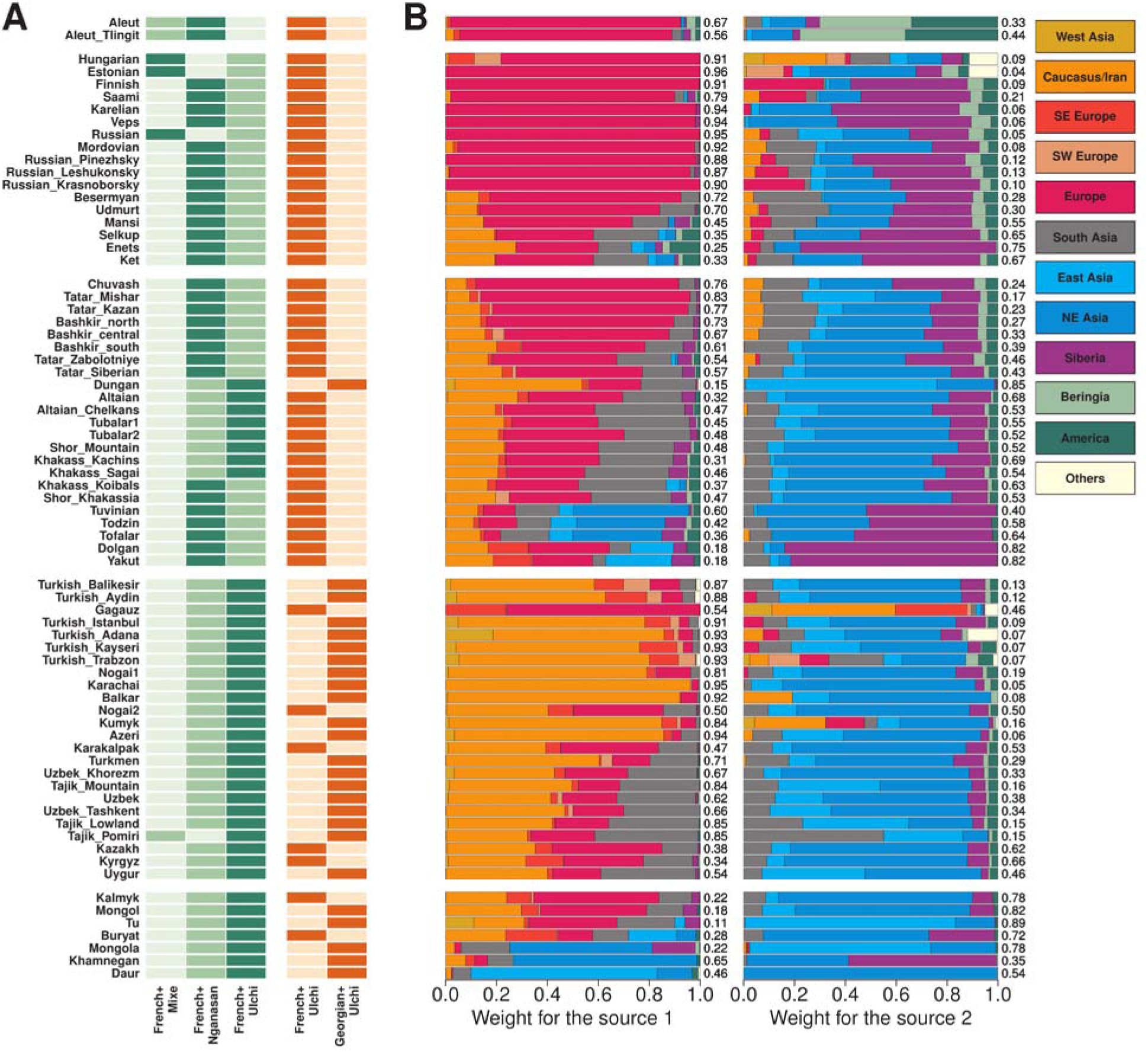
Characterization of the western and eastern Eurasian source ancestries in inner Eurasian populations. (A) Admixture *f*_*3*_ values are compared for different eastern Eurasian references (Mixe, Nganasan, Ulchi; left) or western Eurasian ones (French, Georgian; right). For each target group, darker shades mark more negative *f*_*3*_ values. (B) Weights of donor populations in two sources characterizing the main admixture signal (“date 1 PC 1”) in the GLOBETROTTER analysis. We merged 167 donor populations into 12 groups, as listed on the top right side. Target populations are split into five groups: Aleuts, the forest-tundra cline populations, the steppe-forest cline populations, the southern steppe cline populations and the Mongolic-speaking populations, from the top to bottom.

To perform a higher resolution characterization of the admixture landscape, we performed a haplotype-based GLOBETROTTER analysis. We took a “regional” approach, meaning that all 73 recipient groups were modeled as a patchwork of haplotypes from the 167 donor groups but not those from any recipient group. The goal of this approach was to minimize false negative results due to sharing of admixture history between recipient groups. All of 73 recipient groups show a robust signal of admixture: i.e. a correlation of ancestry status shows a distinct pattern of decay over genetic distance in all bootstrap replicates (bootstrap *p* < 0.01 for all 73 targets; Table S4). When the relative contribution of donors, categorized to 12 groups (Table S2), into the two main sources of the admixture signal (“date 1 PC 1”) is considered, we observe a pattern comparable to PCA, ADMIXTURE and *f*_*3*_ results (Figure 4B). The European donors provide a major contribution for the western Eurasian-related source in the forest-tundra and steppe-forest recipients while the Caucasus/Iranian donors do so in the southern steppe recipients. Similarly, Siberian donors make the highest contribution to the eastern Eurasian-related source in the forest-tundra recipients, followed by the steppe-forest and southern steppe ones.

The GLOBETROTTER analysis also provides an estimate of admixture dates, either one- or two-date estimates, depending on the best model of admixture (Figure S5 and Table S4). We obtain a mean admixture date estimate of 24.3 generations for the steppe-forest and southern steppe cline populations, ranging from 10.7 to 38.1 generations (309 to 1104 years ago, using 29 years per generation^64^). These young dates do not change much even when taking the older dates from the two-date model, as here we obtain a mean of 29.8 generations ranging from 10.7 to 68.1 generations (310 to 1975 years ago). The forest-tundra cline groups have older estimates with a mean of 40.1 generations and a range of 6.8-55.2 generations (197 to 1601 years ago). All but two groups have an estimate older than the steppe mean of 29.8 generations. Estimates of admixture dates using ALDER result in similar values (Figure S5). The admixture dates of the steppe populations are consistent with previous estimates using similar methodologies,^13^ but much younger than expected if they had been driven by admixtures in the Late Bronze and Iron Ages.^8; 38^

### The Eneolithic Botai gene pool provides a glimpse of a lost prehistoric cline

The Eneolithic Botai individuals are closer to each other in the PC space than to any other ancient or present-day individual, and are in proximity to the upper Paleolithic Siberians from the Mal’ta (MA-1) or Afontova Gora (AG3) archaeological sites (Figure 2). Consistent with this, Botai has the highest outgroup *f*_*3*_ statistic with AG3 and other upper Paleolithic Siberians, as well as with the Mesolithic eastern European hunter-gathers from Karelia and Samara (“EHG”) (Figure S6A). East Asians (EAS) are more closely related to Botai than to AG3 as shown by significantly positive *f*_*4*_ symmetry statistics in the form of *f*_*4*_(Mbuti, EAS; AG3, Botai), suggesting East Asian gene flow into Botai (Figure S6B).

We estimated the proportion of East Asian ancestry in Botai using qpAdm. The two-way admixture model of AG3+Korean provides a good fit to Botai with 17.3% East Asian contribution (*Χ*^2^ *p* = 0.286; Table S5), while the models EHG+EAS do not fit (*Χ*^2^ *p* ≤ 1.44×10^−7^). However, we find that Botai harbors an extra affinity with Mesolithic western European hunter-gatherers (“WHG”) unexplained by this model: *f*_*4*_(Mbuti, WHG; AG3+Korean, Botai) is significantly positive in a plausible range of the ancestry proportions (+ 3.0 to + 4.2 SE for 77.7-87.7% AG3 ancestry, mean ± 2 SE; Figure S7). We still obtain a reasonable fit for the same model when we add WHG to the outgroups (*Χ*^2^ *p* = 0.089; Table S5), but adding EHG as an additional source slightly increases model fit with a similar amount of contribution from the East Asian source (*Χ*^2^ *p* = 0.016; 17.3±2.2% East Asian contribution; Table S5).

A graph-based admixture modeling using qpGraph provides similar results: the best two-way admixture model for Botai added to a scaffold graph composed of Mbuti, Onge, Ami, AG3, WHG, and EHG still shows an unexplained affinity between WHG and Botai, and adding an admixture edge from EHG-related branches substantially improves the model fit (Figure S8). Thus, we conclude that the ANE-related ancestry in Botai is intermediate between EHG and AG3, which corresponds to its intermediate geographic position. This suggests a genetic cline of decreasing ANE-related ancestry stretching from AG3 in Siberia to WHG in Western Europe. A substantial East Asian contribution into Botai make them offset from the WHG-ANE cline. A strong genetic affinity between Botai and the Middle Bronze Age Okunevo individuals in the Altai-Sayan region also suggests a wide geographic and temporal distribution of Botai-related ancestry in central Eurasia (Figure S6C).

The Y-chromosome of the male Botai individual (TU45) belongs to the haplogroup R1b (Table S6). However, it falls into neither a predominant European branch R1b-L51^65^ nor into a R1b-GG400 branch found in Yamnaya individuals.^66^ Thus, phylogenetically this Botai individual should belong to the R1b-M73 branch which is frequent in the Eurasian steppe (Figure S9). This branch was also found in Mesolithic samples from Latvia^67^ as well as in numerous modern southern Siberian and Central Asian groups.

### Admixture modeling of contemporary inner Eurasians shows multiple gene flows producing new genetic clines overwriting the ancient ones

Our results show that contemporary inner Eurasians form genetic clines distinct from the ancient WHG-ANE cline, from which a majority of the Botai ancestry is derived. To see if this ancient cline of “ANE” ancestry left any legacy in the genetic structure of inner Eurasians, we performed admixture modeling of populations from the Altai-Sayan region and those belonging to the forest-tundra cline. Specifically, we investigated if an additional contribution from ANE-related ancestry is required to explain their gene pools beyond a simple mixture model of contemporary eastern Eurasians and ancient western Eurasian populations.

Contemporary Altai-Sayan populations are effectively modeled as a two-way mixture of ancient populations from the region with WSH ancestry and contemporary eastern Eurasians, either Afanasievo+Ulchi or Sintashta+Nganasan (*χ*^2^ *p* ≥ 0.05 for 8 / 12 and 5 / 12 Altai-Sayan groups, respectively; Table S7). Among the ancient groups, Sintashta+EAS generally fits Andronovo individuals well with a small eastern Eurasian contribution (6.4±1.4% for estimate ± 1 SE with Nganasans), while later Karasuk or Iron Age individuals from the Altai are modeled better with the older Afanasievo as their WSH-related source (Table S7). If the pre-Bronze Age populations of the Altai-Sayan region were related to either Botai in the west or the Upper Paleolithic Siberians in the east, these results suggest that these pre-Bronze Age populations in southern Siberia did not leave a substantial genetic legacy in the present-day populations in the region. The Okunevo individuals are the only case that WSH+EAS mixture cannot explain (*χ*^2^ *p* ≤ 3.85×10^−4^); similar to Botai, a model of AG3+EAS provides a good fit (*χ*^2^ *p* = 0.396 for AG3+Korean; Table S5).

For the forest-tundra cline populations, for which currently no relevant Holocene ancient genomes are available, we took a more generalized approach of using proxies for contemporary Europeans: WHG, WSH (represented by “Yamnaya_Samara”), and early Neolithic European farmers (EEF; represented by “LBK_EN”; Table S2). Adding Nganasans as the fourth reference, we find that most Uralic-speaking populations in Europe (i.e. west of the Urals) and Russians are well modeled by this four-way admixture model (*χ* ^2^ *p* ≥ 0.05 for all but three groups; Figure 5 and Table S8). Nganasan-related ancestry substantially contributes to their gene pools and cannot be removed from the model without a significant decrease in model fit (4.7% to 29.1% contribution; *χ*^2^ *p* ≤ 1.12×10^−8^; Table S8). The ratio of contributions from three European references varies from group to group, probably reflecting genetic exchange with neighboring non-Uralic groups. For example, Saami from northern Fennoscandia contain a higher WHG and lower WSH contribution (16.1% and 41.3%, respectively) than Udmurts or Besermyans from the Volga river region do (4.9-6.6% and 50.7-53.2%, respectively), while the three groups have similar amounts of Nganasan-related ancestry (25.5-29.1%).

**Figure 5.**
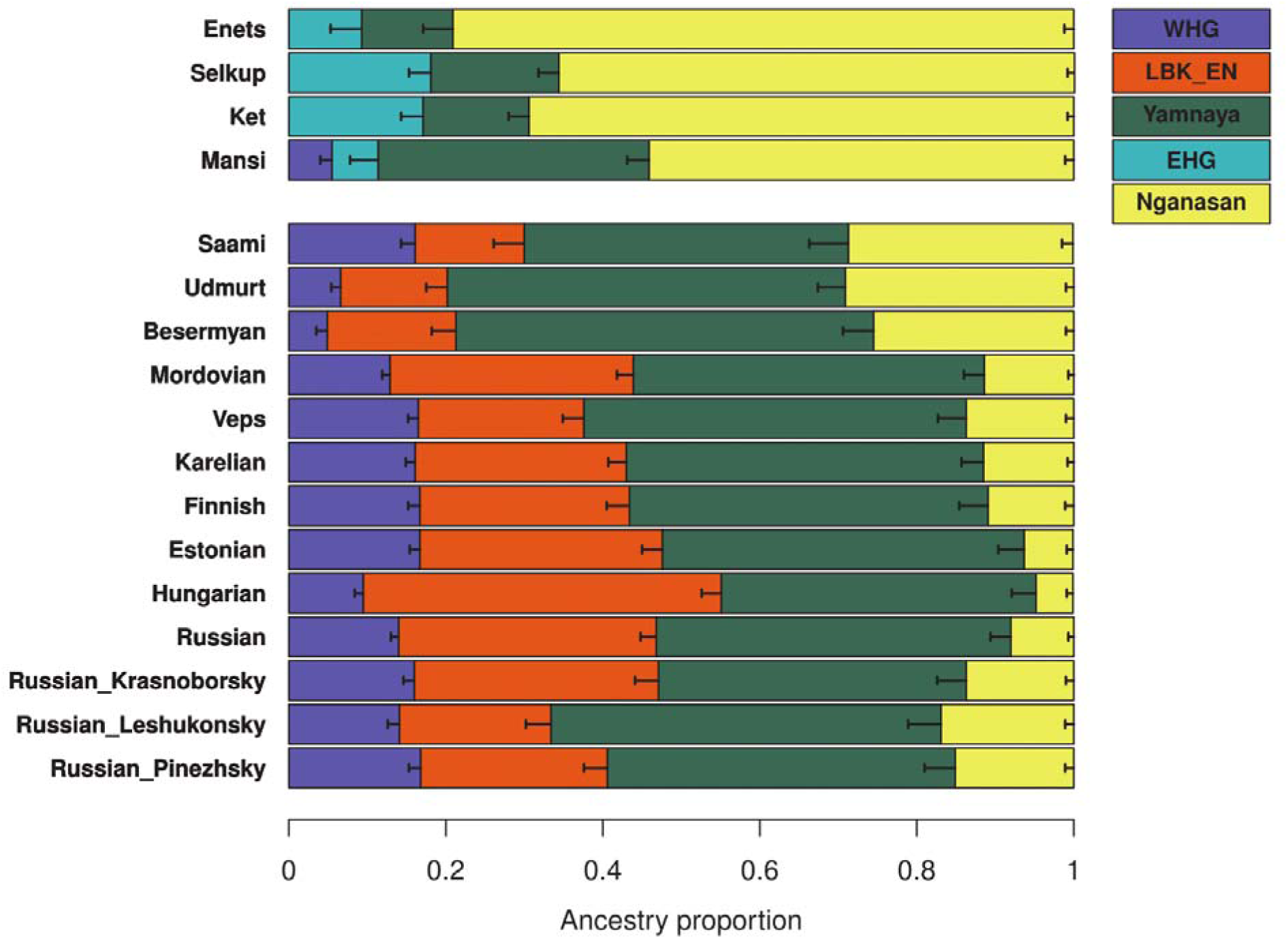
qpAdm-based admixture models for the forest-tundra cline populations. For populations to the east of the Urals (Enets, Selkups, Kets, and Mansi), EHG + Yamnaya + Nganasan provides a good fit, except for Mansi, for which adding WHG significantly increases the model fit. For the rest of the groups, WHG + LBK_EN + Yamnaya + Nganasan in general provides a good fit. 5 cM jackknifing standard errors are marked by the horizontal bar. Details of the model information are presented in Table S8.

For the four forest-tundra cline groups east of the Urals (Enets, Selkups, Kets and Mansi), the above four-way model estimates negative contribution from EEF (< −1.6%). Replacing EEF with EHG, one of the top *f*_*3*_ references for these groups, we obtain well-fitted models with a small WHG contribution (*χ* ^2^ *p* ≥ 0.253; −1.0% to 5.5% WHG contributions). The three-way model excluding WHG shows a good fit for Enets, Selkups and Kets (*χ* ^2^ *p* ≥ 0.098; Figure 5). Simpler models without either EHG or WSH ancestry do not fit (*χ* ^2^ *p* ≤ 0.019 and 0.003, respectively), suggesting a legacy of the ancient WHG-ANE cline.

### The Caucasus Mountains form a barrier to gene flow

When the Altai-Sayan Mountains are often considered as a crossroad of migrations and mark the eastern boundary of the western Eurasian steppe, the Caucasus area plays a similar role for the western end of the steppe. To explore the genetic structure of populations of the Caucasus region, we first performed a PCA of western Eurasians including Caucasus populations (Figure S10). Consistent with previous studies,^4^ Caucasus populations are clustered on the PC space in the vicinity of West Asians further in the south but far from eastern Europeans. The genetic structure within the Caucasus is less pronounced but still evident: populations from the North and South Caucasus, geographically divided by the Greater Caucasus ridge, also show a genetic differentiation. North Caucasus populations show a further subdivision into northwest and northeast groups.

By applying EEMS to the Caucasus region, we identify a strong barrier to gene flow separating North and South Caucasus populations (Figure 6). This genetic barrier coincides with the Greater Caucasus mountain ridge even to small scale: a weaker barrier in the middle, overlapping with Ossetia, matches well with the region where the ridge also becomes narrow. We also observe weak barriers running in the north-south direction that separate northeastern populations from northwestern ones. Together with PCA, EEMS results suggest that the Caucasus Mountains have posed a strong barrier to human migration.

**Figure 6.**
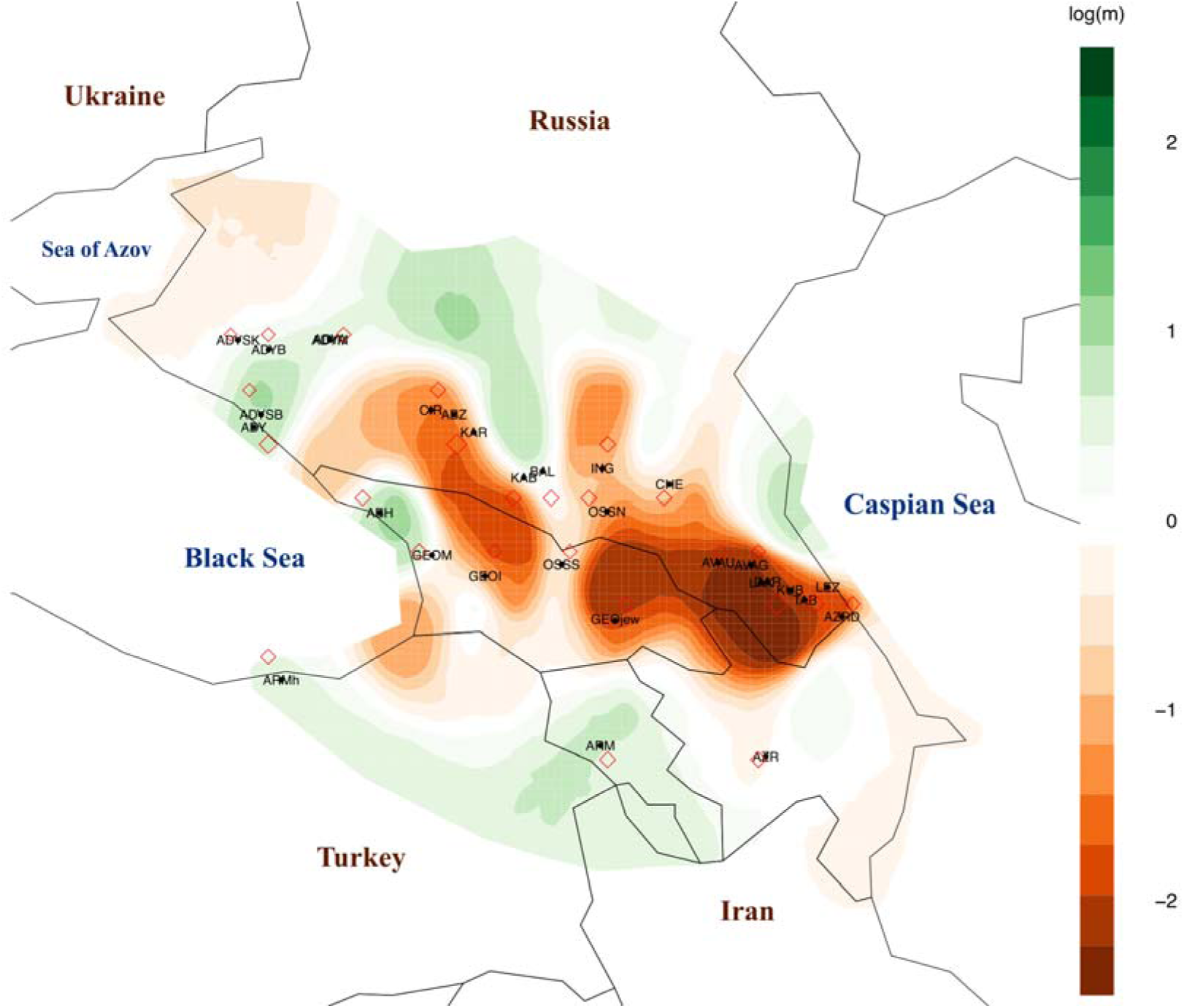
The Greater Caucasus mountain ridge as a barrier to genetic exchange. Barriers (brown) and conduits (green) of gene flow around the Caucasus region are estimated by the EEMSp rogram. Red diamonds show the location of vertices to which groups are assigned. A strong barrier to gene flow overlaps with the Greater Caucasus mountain ridge reflecting the genetic differentiation between populations of the north and south of the Caucasus. The barrier becomes considerably weaker in the middle where present-day Ossetians live.

We quantified the genetic difference within Caucasus populations using *f*_*4*_ statistics of the form *f*_*4*_(Mbuti, X; Caucasus_1_, Caucasus_2_) against world-wide populations outside the Caucasus (“X”). We find many significant *f*_*4*_ statistics suggesting that gene flows from exogenous gene pools have been involved in the development of the population structure of the Caucasus (Figure S11). Compared to both northwest and northeast groups, South Caucasians show extra affinity to Near Eastern populations, such as Neolithic Levantines and Anatolians (“Levant_N” and “Anatolia_N”, respectively; Table S2). In turn, North Caucasus populations have extra affinity with populations of the steppe and broadly of eastern Eurasia. Northeast Caucasians, for example Laks and Lezgins, show the strongest signals with ANE- and WSH-related ancient groups, with MA-1, AG3, Botai and EHG at the top. Northwest Caucasians (e.g. Adygei and Ossetians) are closer to East Asians than Northeast or South Caucasians are. We speculate that these results may suggest at least two layers of gene flow into the North Caucasus region: an older layer related to the ANE- or WSH-related ancestries and the younger layer related to East Asians. The former may have involved an interaction with Iron Age nomads, such as Scythians or Sarmatians. The latter most strongly affected Northwest Caucasians and might be related to historical movements of Turkic populations with some East Asian ancestry into the Caucasus. The genetic legacy of this movement is obvious for the Nogais that are scattered along PC1 between the rest of Caucasus populations and Central Asians (Figures S1-S2).

To explicitly model and quantify the steppe-related gene flows into the Caucasus, we performed a qpAdm-based admixture modeling of 22 Caucasus populations. For 7 of 22 Caucasus populations, a two-way admixture model using Armenians and an ancient Scythian individual^31^ is sufficient (*χ ^2^ p* ≥ 0.05; Table S9). Except for Georgians from the South Caucasus (6.8% contribution from Scythians), all the other groups have a substantial contribution from Scythians (38.0-50.6%). When we add Nanais as the third reference to model potential gene flow from Eastern Eurasians, most of the Caucasus populations are consistent with the model: 15 of 22 Caucasus populations with *χ*^2^ *p* ≥ 0.05 and another three with *χ*^2^ *p* ≥ 0.01 (Table S9). 9 of the 15 groups are adequately modeled by the three references but not by the two: they indeed have positive admixture coefficients for Nanais. Except for Nogais (19.8% for Nogai1 and 48.0% for Nogai2), the other seven groups have only a small amount of East Asian ancestry that is prominent neither in PCA nor in ADMIXTURE (2.7-5.1%; Table S9).

## Discussion

In this study, we analyzed newly reported genome-wide variation data of indigenous people from inner Eurasia, providing a dense representation for human genetic diversity in this vast region. Our finding of inner Eurasian populations being structured into three distinct clines shows a striking correlation between genes and geography (Figures 1-2). The genetic grouping of samples into three clines with gaps in between (Figure 2) corresponds with the fact that samples tend to group into the same clines on the geographic map (Figure 1) with lower density of studied populations in between. However, this non-uniformity of sampling results from the non-uniformity in the density of (language-defined) ethnic groups. Moreover, the reality of the clines was confirmed by the barrier and *f*_*4*_ analyses. The steppe cline populations derive their eastern Eurasian ancestry from a gene pool similar to contemporary Tungusic speakers from the Amur river basin (Figures 2 and 4), thus suggesting a genetic connection among the speakers of languages belonging to the Altaic macrofamily (Turkic, Mongolic and Tungusic families). Based on our results as well as early Neolithic genomes from the Russian Far East,^37^ we speculate that such a gene pool may represent the genetic profile of prehistoric hunter-gatherers in the Amur river basin. On the other hand, a distinct Nganasan-related eastern Eurasian ancestry in the forest-tundra cline suggests a substantial separation between these two eastern ancestries. Nganasans have high genetic affinity with prehistoric individuals with the “ANE” ancestry in North Eurasia, such as the Upper Paleolithic Siberians or the Mesolithic EHG, which is exceeded only by Native Americans and by Beringians among eastern Eurasians (Figure S12). Also, Northeast Asians are closer to Nganasans than they are to either Beringians or Native Americans, and the ANE affinity in East Asians is correlated well with their affinity with Nganasans (Figure S13). We hypothesize that Nganasans may be relatively isolated descendants of a prehistoric Siberian gene pool, which formed modern Northeast Asians by mixing with populations related to the Neolithic Northeast Asians.^37^

The Botai genomes provide a critical snapshot of the genetic profile of pre-Bronze Age steppe populations. Our admixture modeling positions Botai primarily on an ancient genetic cline of the pre-Neolithic western Eurasian hunter-gatherers: stretching from the post-Ice Age western European hunter-gatherers (e.g. WHG) to EHG in Karelia and Samara to the Upper Paleolithic southern Siberians (e.g. AG3). Botai’s position on this cline, between EHG and AG3, fits well with their geographic location and suggests that ANE-related ancestry in the East did have a lingering genetic impact on Holocene Siberian and Central Asian populations at least till the time of Botai. A recent study reports 6,000 to 8,000 year old genomes from a region slightly north of Botai, whose genetic profiles are similar to our Botai individuals.^68^ This ancient cline in Altai-Sayan region has now largely been overwritten by waves of genetic admixtures. Starting from the Eneolithic Afanasievo culture, multiple migrations from the Pontic-Caspian steppe to the east have significantly changed the western Eurasian ancestry during the Bronze Age.^7; 8^ Our admixture modeling finds that no contemporary population in the Altai-Sayan region is required to have additional ANE ancestry beyond what the mixture model of Bronze Age steppe plus modern Eastern Eurasians can explain (Table S7). The most recent clear connection with the Botai ancestry can be found in the Middle Bronze Age Okunevo individuals (Figure S6C). In contrast, additional EHG-related ancestry is required to explain the forest-tundra populations to the east of the Urals (Figure 5 and Table S8). Their multi-way mixture model may in fact portrait a prehistoric two-way mixture of a WSH population and a hypothetical eastern Eurasian one that has an ANE-related contribution higher than that in Nganasans. Botai and Okunevo individuals prove the existence of such ANE ancestry-rich populations. Pre-Bronze Age genomes from Siberia will be critical for testing this hypothesis.

The study of ancient genomes from inner Eurasia will be extremely important for going forward. Inner Eurasia has functioned as a conduit for human migration and cultural transfer since the first appearance of modern humans in this region. As a result, we observe deep sharing of genes between western and eastern Eurasian populations in multiple layers: the Pleistocene ANE ancestry in Mesolithic EHG and contemporary Native Americans, Bronze Age steppe ancestry from Europe to Mongolia, and Nganasan-related ancestry extending from western Siberia into Eastern Europe. More recent historical migrations, such as the westward expansions of Turkic and Mongolic groups, further complicate genomic signatures of admixture and have overwritten those from older events. Ancient genomes of Iron Age steppe individuals, already showing signatures of west-east admixture in the 5^th^ to 2^nd^ century BCE,^38^ provide further direct evidence for the hidden old layers of admixture, which is often difficult to appreciate from present-day populations as shown in our finding of a discrepancy between the estimates of admixture dates from contemporary individuals and those from ancient genomes.

## Supplemental Data

Supplemental Data include 13 figures and 9 tables.

## Acknowledgements

We thank Iain Mathieson and Iosif Lazaridis for their helpful comments. The research leading to these results has received funding from the Max Planck Society, the Max Planck Society Donation Award and the European Research Council (ERC) under the European Union’s Horizon 2020 research and innovation programme (grant agreement No 646612 granted to M.R.). Analysis of the Caucasus dataset was supported by RFBR grant 16-06-00364 and analysis of the Far East dataset was supported by Russian Scientific Fund project 17-14-01345. D.R. was supported by the U.S. National Science Foundation HOMINID grant BCS-1032255, the U.S. National Institutes of Health grant GM100233, by an Allen Discovery Center grant, and is an investigator of the Howard Hughes Medical Institute. P.F. was supported by IRP projects of the University of Ostrava and by the Czech Ministry of Education, Youth and Sports (project OPVVV 16_019/0000759). C.C.W. was funded by Nanqiang Outstanding Young Talents Program of Xiamen University and the Fundamental Research Funds for the Central Universities. M.Z. has been funded by research grants from the MES RK No. AP05134955 and No. 0114RK00492.

## Web Resources

IMPUTE version 2 (IMPUTE2), https://mathgen.stats.ox.ac.uk/impute/1000GP_Phase3.html

International Society of Genetic Genealogy (ISOGG), http://www.isogg.org pileupCaller, https://github.com/stschiff/sequenceTools

Sequence Read Archive (SRA), https://www.ncbi.nlm.nih.gov/sra

YFULL™.com, http://www.yfull.com

## Accession Numbers

Genome-wide sequence data of two Botai individuals (BAM format) are available at the Sequence Read Archive under the accession number PRJNA470593. Array genotype data will be made available through the Reich Lab and MPI-SHH webpages upon the publication of the manuscript.

